# Thalamocortical transmission of visual information in awake mice involves phase synchronization and spike synchrony at high gamma frequencies

**DOI:** 10.1101/170167

**Authors:** Samuel S. McAfee, Yu Liu, Mukesh Dhamala, Detlef H. Heck

## Abstract

Synchronization of neuronal spike activity is thought to play a key role in the transmission of information for sensory processing in the brain, and this synchronization is influenced by oscillatory population activity occurring in multiple frequency ranges at multiple stages of sensory pathways. In the neocortex, gamma frequency oscillations appear to play an important role in synchronizing neuronal ensembles and allowing for selective communication between regions, yet relatively little is known about whether gamma oscillations facilitate transmission of sensory information from thalamus to cortex. Here, we investigate the role of gamma oscillations in promoting synchronous spike activity between the visual thalamus (dLGN) and primary visual cortex (V1) in awake mice, a model sensory system with prominent gamma oscillations that are modulated by visual input. We demonstrate that transmission of visual information to cortex involves phase-synchronized oscillations in the high gamma range (50-90Hz), with concomitant millisecond-scale synchronization of thalamic and cortical spike activity. Transition from a full-field gray image to a high-contrast checkerboard image caused gamma activity to rapidly increase in amplitude, frequency, and bandwidth, yet the gamma oscillations in dLGN and V1 maintained a consistent phase relationship. High contrast stimulation also caused an increase in amplitude of oscillations in the beta and low gamma range, but those were not associated with synchronous thalamic activity. These results indicate a role for high gamma oscillations in mediating the functional connectivity between thalamic and cortical neurons in the visual pathway, a similar role to beta oscillations in primates.

**Significance statement:** The mechanisms by which neurons selectively communicate are essential to our understanding of how the brain processes information. Abundant evidence suggests that cortical sensory processing involves the synchronization of high frequency electric field oscillations known as gamma oscillations, which allow groups of neurons to synchronize their spike activity in order to collaboratively process sensory input. Here, we show that oscillations and spikes in the visual thalamocortical pathway of the mouse exhibit synchrony across a broad high gamma frequency range (50-90Hz), suggesting these oscillations play an important role in the relay of visual information to the cortex. This is substantially different from oscillations observed in monkeys, in which gamma is absent in thalamus and beta oscillations support thalamocortical relay.

## Introduction

Sensory processing requires selective and effective communication between neuronal groups within and across brain regions, and this is understood to be facilitated by the appropriate phase alignment of regional oscillations (Bastos, Vezoli et al. 2015). Connected regions can transiently increase the effectiveness of their communication (functional connectivity) by aligning their oscillations so that local neurons are driven to generate output that arrives at target regions at moments of maximal gain (Fries, Nikolic et al. 2007, Womelsdorf, Schoffelen et al. 2007, Fries 2015). This is known as phase synchronization, and it likely underlies both the initial high-fidelity relay of information along a sensory pathway, as well as the recruitment of cell assemblies within a region and the selective communication among them.

In the visual cortex (V1) of a variety of species, oscillatory activity and neuronal firing synchrony are known to be stimulus-dependent, meaning there is a strong correlation of image properties such as spatial frequency and contrast with the power and frequency of cortical gamma oscillations (Henrie and Shapley 2005, Ray and Maunsell 2010, Roberts, Lowet et al. 2013), as well as long-range synchrony between neurons with similar stimulus preferences (Gray, Konig et al. 1989, Nase, Singer et al. 2003, Kohn and Smith 2005, Bachatene, Bharmauria et al. 2016). Inter-areal phase synchronization is most commonly studied as it applies to different cortical areas binding contiguous visual images (Usher and Donnelly 1998, Engel, Fries et al. 2001, Veit, Hakim et al. 2017), but also likely plays a role in the dedicated relay of visual information from the dorsolateral geniculate nucleus of the thalamus (dLGN) to V1 (Castelo-Branco, Neuenschwander et al. 1998, Bastos, Briggs et al. 2014).

Little is known about the role of phase synchronization in thalamocortical transmission in awake animals, and the available evidence suggests there may be considerable differences in oscillation frequencies between species. In primates, experimental evidence suggests thalamocortical transmission is supported by oscillations in the beta frequency range (Bastos, Briggs et al. 2014), with gamma oscillations first being generated in the cortex. In contrast to this, recently published studies in mice suggest that a prominent narrowband oscillation in the high gamma range may underlie visual thalamocortical transmission (Saleem, Lien et al. 2017, Veit, Hakim et al. 2017). This narrowband oscillation originates in the retina in response to luminance of the visual field (Koepsell, Wang et al. 2009), and retinal activity drives coherent oscillations as far downstream as layer 2/3 of V1 (Saleem, Lien et al. 2017, Veit, Hakim et al. 2017). Somewhat problematically for this role however, the introduction of visual contrast dramatically changes the activity in visual cortex, leading to the suppression of gamma amplitude in the narrow band and the emergence of distinct broadband low gamma and high gamma oscillations. Here we asked whether one of these contrast-evoked gamma frequency oscillations is related to thalamocortical input. We hypothesized that either contrast induces emergent cortical gamma oscillations, which lose alignment with narrowband thalamic input, or contrast drives similar broadband thalamic oscillations, which maintain a consistent alignment with those in cortex.

Here we addressed these possibilities by performing dual dLGN-V1 recordings in awake, head-fixed mice presented with zero-contrast and high contrast visual stimuli. We asked whether high-contrast images induce synchronized gamma oscillations in mouse dLGN and V1, and investigated whether V1 oscillations were linked to the occurrence of synchronous spikes thought to reflect effective communication between the two structures. We found that the transition from the full-field gray image to a high-contrast checkerboard image lead to stimulus-locked phase synchronization of high gamma oscillations in V1 and dLGN within the range of 50-90Hz, immediately following the rhythmic occurrence of synchronous spike activity in both regions. These findings suggest that thalamocortical transmission involves a range of coherent high gamma oscillations, and that synchronization of high gamma is likely to constitute a dedicated frequency channel for the thalamocortical relay of visual information in mice.

## Methods

### Animals

Experiments were performed on both Male and Female C57BL/6J mice (>8 weeks old, 18-28 g body weight). Mice were housed in a breeding colony with 12-hour light/dark cycles with ad libitum access to food and water. All mice were housed in the dark for 12 hours prior to recording. All experimental procedures adhered to guidelines approved by the University of Tennessee Health Science Center Animal Care and Use Committee. Principles of laboratory animal care (NIH publication No. 86-23, rev. 1996) were followed.

### Preparation for awake, multisite recordings

Mice were induced for surgery with 3% Isoflurane (Baxter Pharmaceutical Products, Deerfield IL) in oxygen in an incubation chamber, and transferred to a stereotaxic head mount and anesthesia was maintained with 1–2.5% Isoflurane in oxygen through a nose cone. Isoflurane concentration was controlled with a vaporizer (Highland Medical Equipment, CA). The depth of anesthesia was adjusted until the mice failed to show a reflex withdrawal of the hind paw to a strong pinch. Blunt ear bars were used to prevent damaging the eardrums. Core body temperature, measured with a rectal thermometer, was maintained between 36.5 and 38.0 °C with a feedback-controlled heating pad (FHC Inc., Bowdoinham, ME). Surgical techniques were described in detail elsewhere. In brief, a small craniotomy (2 mm x 1 mm) was made above the primary visual cortex and dorsal lateral geniculate nucleus. The exposed but intact dura was covered with Triple Antibiotic (Walgreens, US) to maintain moisture and reduce the risk of infection. A cylindrical plastic chamber (0.45 cm diameter and 4 mm height) was placed over the skull opening and filled with Triple Antibiotic and Kwik-sil epoxy (World Precision Instruments, Sarasota, FL). Three small machine screws (1/8′ dome head, 0.8 mm diameter, 2 mm long, Small Parts, Inc., Miami Lakes, FL) were secured in the skull bone, and a metal head post was mounted anterior to Bregma. The chamber, head post and skull screws were secured in place with super glue and dental acrylic. Mice were injected subcutaneously with 5 mg kg^−1^ analgesic Carprofen (Zoetis Inc.; Kalamazoo, MI) to alleviate pain and 0.25 ml of lactated ringer solution as a fluid supplement twice within the first 24 h of the surgery. Mice were allowed a 3-4 day recovery from surgical preparation before subsequent procedures.

### Electrophysiological recordings

Mice were habituated to head fixation above a freely rotating treadmill for one 30-minute session before recording sessions started. The experimental setup, head-holding device and recording procedures have been described in detail previously(Bryant, Roy et al. 2009). Head fixation was accomplished by tightening a mechanical screw attached to a fixed bar through a threaded hole in the metal head post. Mice were allowed to move and walk freely on the treadmill at all times. The plastic chamber was then cleaned and filled with sterile saline solution.

LFP recordings were accomplished using five hand-made glass insulated tungsten/platinum electrodes (impedance 1.0-5.0 MΩ) guided in the ventral-dorsal direction by a computer-controlled microdrive (MiniMatrix, Thomas Recording, Germany). Stainless steel microdrive guiding tubes served as reference electrodes, and were electrically connected to the brain tissue via the electrolyte solution in the chamber. Guiding tubes were positioned at the surface of the brain above the targeted structures, and the electrodes were slowly advanced through the dura to their approximate target depths. Electrode positions were then finely adjusted during the presentation of visual stimuli in order to preliminarily identify the locations of electrode tips based on established patterns of evoked neuronal activity(Niell and Stryker 2008). Electrode movements were controlled with micrometer resolution and digitally monitored. Local field potentials and spike signals were separated by band pass filtering at 0.1 to 200 Hz and at 200 Hz to 8 kHz, respectively, using a hardware filter amplifier (FA32; Multi Channel Systems). Filtered and amplified voltage signals were digitized and stored on a computer hard disk (16 bit A/D converter; sampling rate, >20 kHz for action potentials, >2kHz for LFPs) using a CED power1401 and Spike2 software (both Cambridge Electronic Design).

Upon completion of each recording session, electrolytic lesions were created in the tissue at the electrode tips in order to confirm recording sites. The electrodes were then retracted, saline was removed from the chamber, triple antibiotic was reapplied, and the mice were temporarily placed back into their home chambers. All mice were euthanized within 8 hours of recording and transcardially perfused with phosphate-buffered saline and 4% paraformaldehyde prior to brain tissue collection.

### Presentation of visual stimuli

Visual stimuli were designed and presented using customized psychtoolbox and Matlab scripts(Brainard 1997, Pelli 1997). Checkerboard images (100% contrast, 0.05 cpd, 100ms duration) were presented on a 22″ computer monitor centered 30° lateral to the animal’s midline and 25 cm away. Stimuli were presented at randomly generated intervals ranging from 2.5 to 4 seconds. The monitor remained gray, with the same luminance as the checkerboard during the inter-stimulus intervals. Precise timing of visual stimuli was co-registered to the LFP by simultaneously recording the voltage from a photodiode placed on the corner of the screen. LFP recordings were taken on the contralateral side to the visual field in which stimuli were presented.

### Analysis of Local Field Potential amplitude and phase

Recorded data were visually inspected and periods that were free of movement artifact or treadmill rotation from locomotion were selected for further analysis. 60Hz contamination of the signal by ambient electrical noise was removed using a hum-removal algorithm in the Spike2 software. LFP data and stimulus times were then exported to Matlab for further analysis.

Frequency components of each LFP signal were extracted using acausal FIR band pass filters. Filters of bandwidth 0.5-5Hz were used in 0.5Hz steps, with filter order equal to the number of samples in 5 cycles for each center frequency. An amplitude time series for each frequency was estimated by applying the Hilbert transform to each band pass filtered signal and taking its absolute value. Changes in amplitude were calculated as a percent difference from the mean values occurring within 1s prior to the stimuli. Instantaneous phase values were determined at each time point as the angular value of the analytic signal. Pairwise-phase consistency (PPC)(Vinck, van Wingerden et al. 2010) values were calculated on phase values across trials for each frequency and latency from the stimulus in 1ms time steps.

### Time resolved LGN-V1 spike synchrony

In order to assess the connection between oscillatory synchrony and neuronal communication between V1 and dLGN neurons, we focused on the occurrence of synchronous firing of multiple units between the regions. Spikes were detected using the offline spike sorting algorithm in Spike2 (CED details), and spike times were exported to Matlab for further analysis.

A strong interest in this study was to determine whether rhythmic occurrences of synchronous spikes could be detected in the recording in a manner that allowed for later decomposition into frequency components. This type of approach would require a conversion of discrete spike events into a continuous function, which allows for interpretation of individual stimulus presentations as well as averaging across them. To accomplish this, we created a continuous time series reflecting spike synchrony by sliding a window across the recording period and measuring the coincidence of V1 and dLGN spikes (e.g. the product of their counts) within the window. We shifted the time of dLGN spikes by 5ms to account for conduction delay, and counted coincidences within a conservative 4ms time window roughly corresponding to the naturally occurring variability in latency of evoked V1 responses (MacLean, Fenstermaker et al. 2006). This window width is also within the interval that synchrony of dLGN outputs evokes maximal V1 responses (Wang, Spencer et al. 2010). The mean of 200 jittered data sets (±2-10ms jitter range, 0.5ms resolution, uniform probability) was then subtracted in order to correct for average synchrony occurring due to increases in spike rate alone, which creates a positive offset in the signal and obscures the content of interest. The jittering method was found to be preferable to using a trial shift or shuffle predictor, as it disrupted only the precise timing of spikes while preserving the approximate latency and specific spike count in each response (Torre, Canova et al. 2016). The function was then smoothed with a 4ms window. This process revealed rhythmicity in the synchronous occurrence of spikes, which could then be the subject of individual trial investigations as well as time-frequency analysis in the same manner as the LFP.

### Correlation of Spike synchrony and Gamma activity

Lastly, we investigated whether synchronously occurring spikes in dLGN and V1 were consistently linked to the occurrence of V1 oscillations at different frequencies. To do this, we used the synchrony function described in the previous subsection to perform a cross correlation analysis with different band pass filtered components of the V1 LFP. Sections corresponding to gray screen and checkerboard viewing were separated before further analysis in order to find differences between the two states.

### Experimental Design and Statistical Analysis

Six mice (2 female, 4 male) underwent successful electrophysiological recording in the current study. Successful recording was defined as having one or more units in each location that responded positively to contrast increase, having at least 250 artifact-free stimulus cycles executed over a 15 minute recording session, and having histological confirmation of recording in dLGN and layer 4 of V1. Twelve mice underwent at least some portion of the aforementioned surgical and experimental procedures but failed to meet these criteria.

All statistical analyses were performed in Matlab. Statistical assessment of correlations between LFP oscillations and spikes was performed using non-parametric Monte Carlo approaches with surrogate data. For assessment of the correlation between PPC and V1 oscillation amplitude, surrogate values were generated for each evaluated frequency by randomly shifting PPC results across time in a circular manner with respect to the mean LFP amplitude of the same frequency. 200 surrogate values were generated for each frequency for each animal. The surrogate coefficients were then ranked, and the 95^th^ and 5^th^ percentile coefficients for each frequency were used as a statistical threshold. Averaging of correlation values for graphical representation was performed following a Fischer transformation of the coefficients, and then the inverse on the mean. R^2^ values for the peak V1 high gamma frequencies were created by taking the square of the raw coefficient, and averaged across animals.

For analysis of the relationship between LGN-V1 spike synchrony and the V1 LFP, we developed a Monte Carlo hypothesis test for the precision of spike times and their temporal relationship to V1 oscillations in the high gamma range (50-90Hz). For this approach, real correlations were tested against a ranked distribution of surrogate correlation values between the V1 LFP and jittered spike synchrony waveforms. In order to conservatively test for a relationship between thalamocortical spike synchrony and V1 gamma without bias caused by the phase locking of V1 spikes to V1 gamma, only dLGN spike times were jittered in the generation of surrogates. Correlation values above the 95^th^ percentile or below the 5^th^ percentile were considered significant.

## Results

### Visual stimulus-driven gamma oscillations in dLGN and layer 4 of V1

As in other studies, we observed high amplitude gamma oscillations in layer 4 of V1 in response to the presentation of a high-contrast checkerboard image (Niell and Stryker 2008, Saleem, Lien et al. 2017, Veit, Hakim et al. 2017) as well as “narrowband” gamma oscillations during periods of viewing the gray screen (Saleem, Lien et al. 2017) (Figure 1). By comparing contrast-evoked oscillations to the amplitudes of corresponding oscillations in the gray screen condition, we were able to identify activity changes associated with contrast onset, and characterize the rapid transition between the two states (Figure 1C-G). V1 showed a dramatic amplitude increase in the high beta/low gamma (20-35Hz) and high gamma range (50-90Hz), the latter of which was higher in frequency and had broader bandwidth than the previously reported narrowband gamma (Saleem, Lien et al. 2017, Veit, Hakim et al. 2017). The dLGN LFP showed no such increase in beta or low gamma but did show increased high gamma amplitude, with frequencies overlapping with gamma oscillations in V1. Time-frequency analysis revealed that high gamma oscillations arose in dLGN before V1, consistent with a causal role.

**Figure 1:**
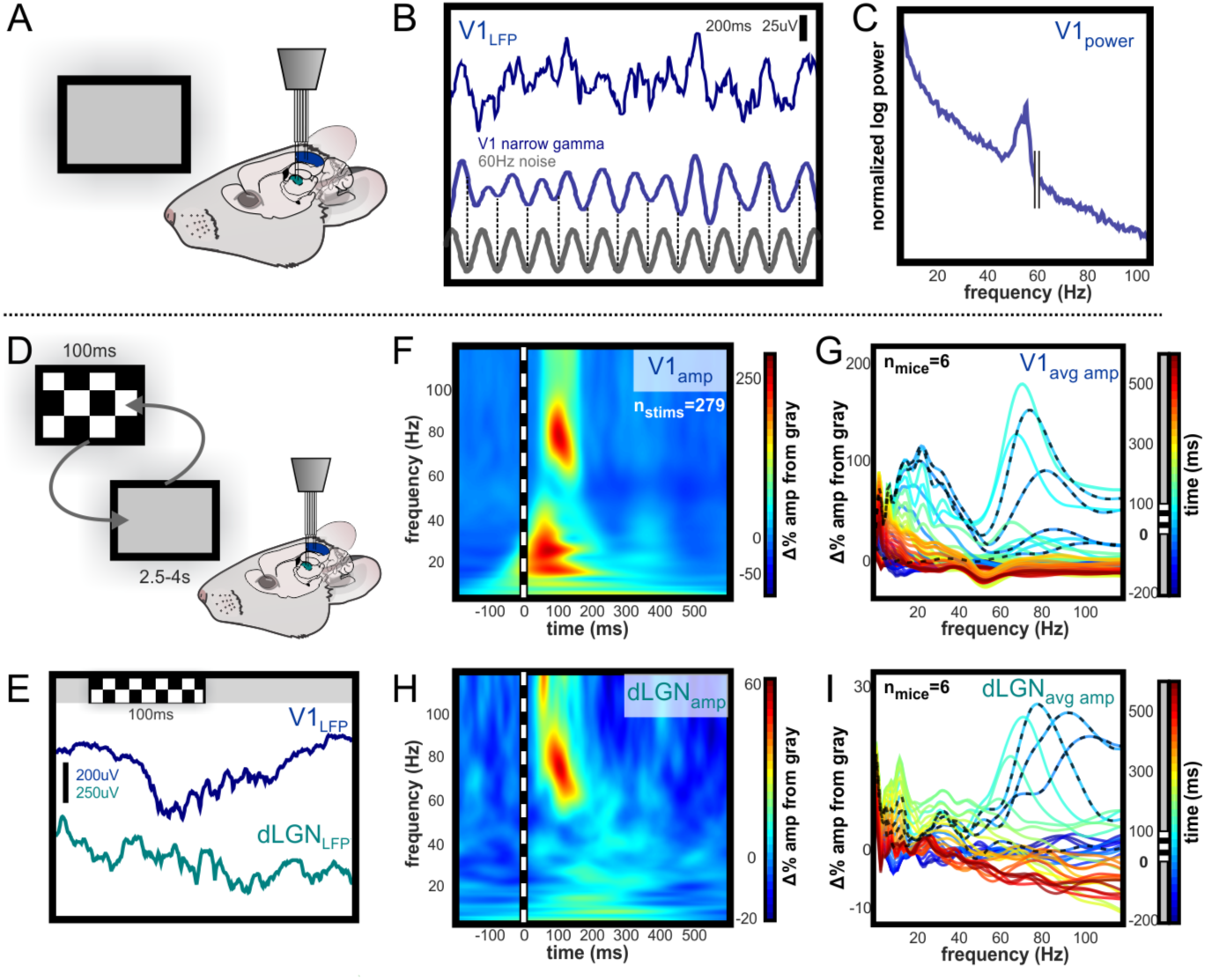
Visual-evoked narrowband and broadband high gamma oscillations in the visual system. Independently guided electrodes were used to record LFP and spikes from L4 of V1 and dLGN during visual stimulation. A: Illustration of experimental setup during baseline recording of activity evoked by viewing of the gray computer screen. B: Example of 200ms of raw narrowband gamma in L4 of V1 evoked by luminance of a gray computer screen. The frequency is between 50 and 60Hz, but ambient 60Hz electrical noise is not linked to this oscillation, as illustrated by the phase shifts near the beginning and end of trace. C: Spectral power of narrowband gamma oscillation in V1. Peak in spectral power is apparent below 60Hz. D: Illustration of experimental setup during randomized presentation of high contrast checkerboard image E: Brief high-contrast checkerboard stimuli were presented and changes in oscillatory activity were observed in V1 and dLGN. Contrast-evoked oscillations are higher in amplitude and wider in bandwidth than narrowband gamma in V1, and are also visible in dLGN. F: Example time-frequency analysis of contrast-evoked LFP oscillations in V1, normalized to amplitude during gray screen viewing. 100ms checkerboard image evokes high amplitude oscillations in distinct low and high gamma frequency ranges. G: Group mean change in oscillation amplitude over time relative to LFP during gray screen viewing. Each line represents amplitude within a 25ms time bin. Peak amplitude change occurs 100-125ms after stimulus onset in the 50-90Hz range. Narrowband gamma remains decreased immediately after checkerboard offset. H,I: Same as D,E for LFP in dLGN. Low gamma oscillation amplitude is not modulated in dLGN, and high gamma oscillations arise in LGN before V1.

### Phase synchronization and amplitude of LFP oscillations

Gamma oscillations in the cortex are most commonly generated and maintained by local cycles of inhibitory activity (Fries, Nikolic et al. 2007, Whittington, Cunningham et al. 2011, Buzsaki and Wang 2012), however, narrowband high gamma activity in V1 was shown to be driven by excitatory input from dLGN, even when cortex was silenced (Saleem, Lien et al. 2017). To explore the hypothesis that broadband high gamma oscillations occurring in V1 were driven by rhythms in dLGN, we investigated which V1 oscillations concurrently increased in amplitude while phase coupling with frequency-matched dLGN oscillations. In order to do this, we performed a correlation analysis on the average time course of phase synchronization (PPC) and the amplitude of oscillations in V1, a similar approach to what was developed for a comparable investigation in primates (Bastos, Briggs et al.). This revealed that V1 high gamma amplitude co-varied with the strength of phase synchronization with dLGN (significant in 6 of 6 animals), with dLGN-V1 phase synchrony accounting for 66.9% of the amplitude in the peak V1 high gamma frequency across animals. A consistent inverse pattern was found in the alpha range, such that alpha oscillations in V1 became uncoupled from those in dLGN as contrast drove alpha amplitude increases in V1 (significant in 6 of 6 animals; Figure 2C-D).

**Figure 2:**
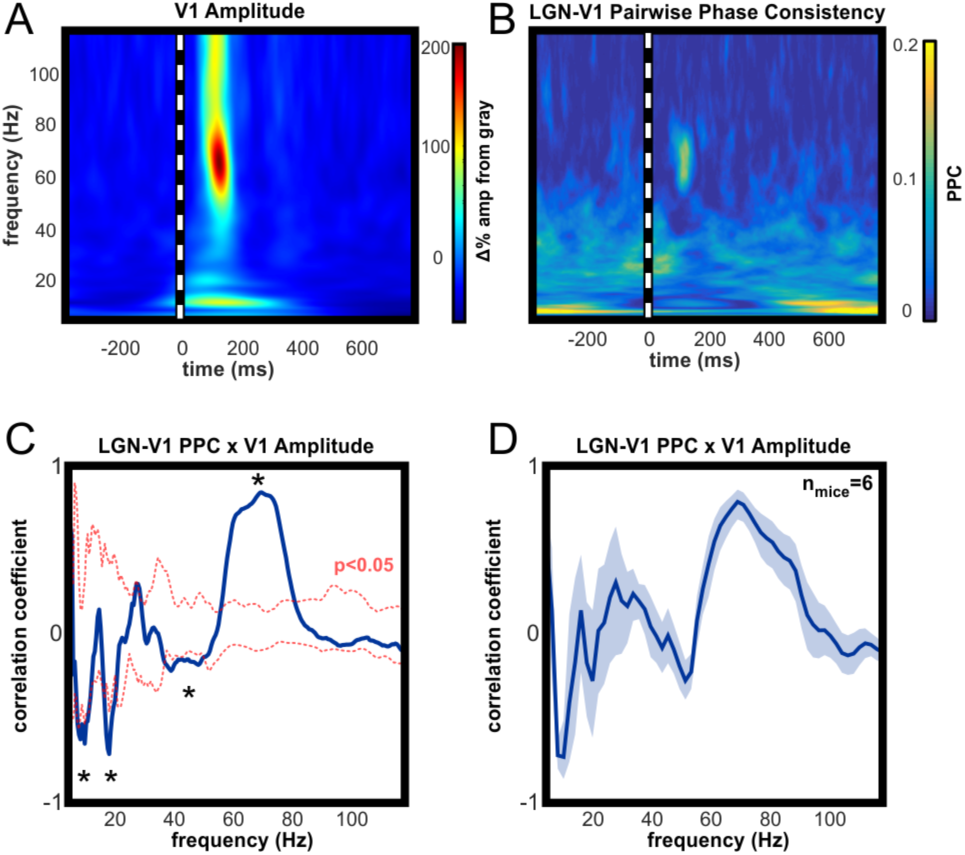
Amplitude of high gamma oscillations is strongly linked to simultaneous thalamic high gamma activity. A: Example time-frequency analysis of V1 LFP, aligned on the presentation of 100ms full-contrast checkerboard image. B: Example time-resolved pairwise phase consistency analysis of oscillation phase synchronization between dLGN and V1. PPC is increased in the high gamma frequency range following checkerboard stimulus. C: Correlation analysis of A and B to identify V1 oscillations whose amplitude is dependent on synchronous thalamic oscillations across stimulus conditions. High gamma in V1 is strongly correlated with synchronous oscillations in dLGN, and strongly anti-correlated with synchrony in the alpha, beta and low gamma ranges. D: Group mean results (±SEM) for correlation analysis in C. All animals had significant correlations within the 50-90Hz high gamma frequency range and significant inverse correlations in the alpha frequency range. At the peak high gamma frequency in V1, dLGN-V1 phase synchronization accounted for an average of 66.9% of the amplitude modulation, across animals.

### dLGN-V1 spike synchrony

Sudden changes in visual stimuli are often characterized by rapid burst-firing of neurons in the dLGN (Grubb and Thompson 2005), which we observed in response to the checkerboard image. Onset of contrast-induced spiking in V1 units typically occurred at the offset of bursting in dLGN units, and these initial spikes were synchronized with those in dLGN above mean probability based on spike rate alone (Figure 3).

**Figure 3:**
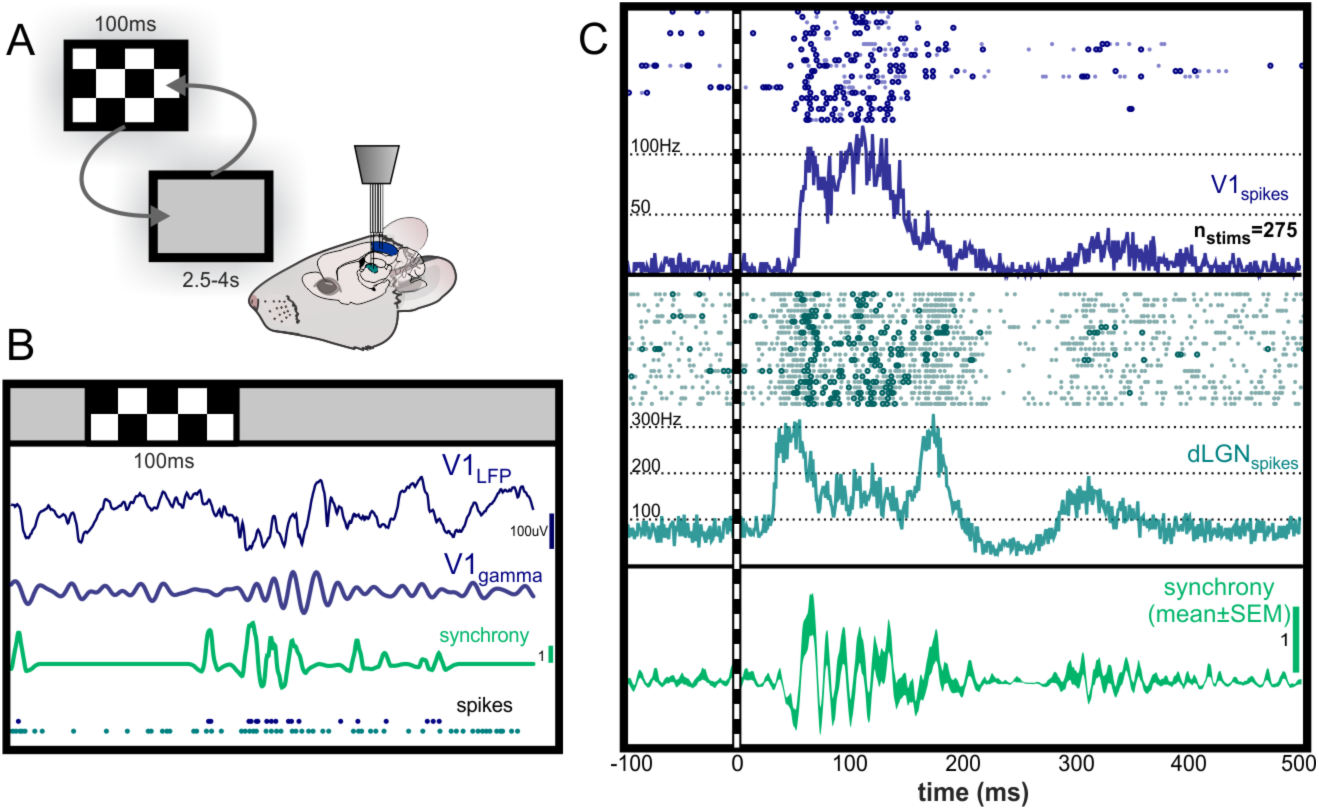
Time-resolved estimation of spike synchrony is rhythmic in the high gamma range. A: Illustration of experimental setup during randomized presentation of high contrast checkerboard image. B: Single trial example of spike synchrony following checkerboard stimulus. Green trace represents time-resolved spike synchrony function, calculated from V1 and dLGN spike times. Synchronous bursts of spikes are observed following the checkerboard image. Scale bar shows height of 1 coincidence above mean probability. C: Example raster for V1 and dLGN spikes occurring in 20 consecutive trials, and peri-stimulus histogram over all trials. Circles in raster highlight spikes occurring synchronously above mean probability. After initial dLGN burst and V1 response, dLGN spikes appear to be modulated at an approximate 68Hz rhythm. Bottom panel shows synchrony function averaged over trials to reveal rhythmicity of synchronous spikes locked to the stimulus onset.

Occurrence of synchronized spikes began after initial dLGN bursts, but before the onset of high amplitude broadband gamma in V1. Time-frequency analysis of the spike synchrony revealed peak rhythmicity in the high gamma range, which overlapped with the frequency of evoked gamma oscillations in V1 (Figure 4). The duration of this rhythmic interaction overlapped with the emergence of high gamma in the V1 LFP.

**Figure 4:**
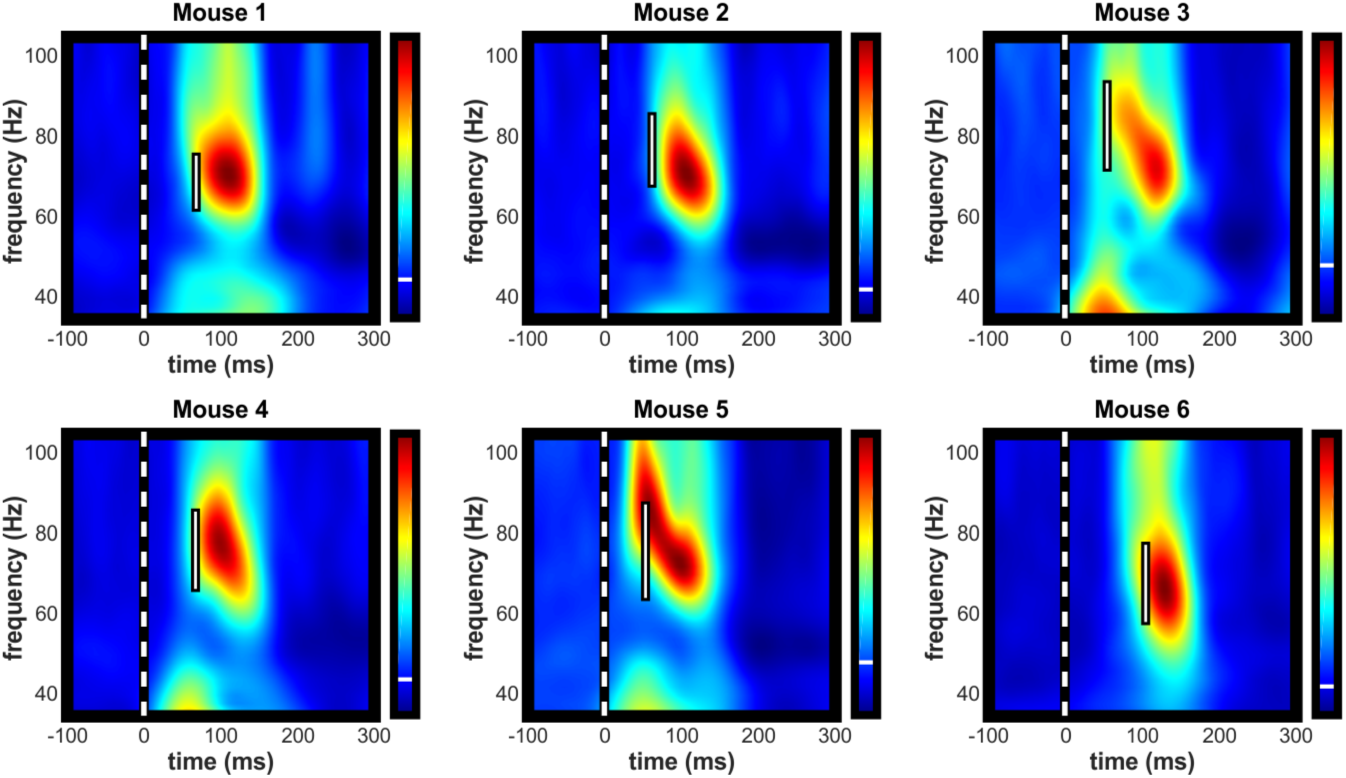
Amplitude increase of high gamma oscillations in V1 LFP follows synchronization of spike activity in the same frequency range. Individual examples of contrast-evoked gamma oscillations with latency and bandwidth of peak spike synchrony superimposed. White bar spans the bandwidth of 95% maximal amplitude of rhythmic spike synchrony. All color axes are normalized to maximum amplitude increase and decrease with white line indicating zero amplitude change.

Broad stimulus-evoked spike synchrony between visually-responsive neurons has been reported in mouse cortex at recording sites separated by a distance of 350um (Nase, Singer et al. 2003) to 500um (Veit, Hakim et al. 2017). Based on estimations from published images of intact thalamocortical projections *in vitro* (MacLean, Fenstermaker et al. 2006), the length of visual thalamocortical axons, which project to the cortex via the white matter tract along the dorsolateral edge of the hippocampal formation, is within the range of 5000-6500um. This is a far greater length than the broadest width of the V1, thus the precise synchrony we observe between thalamic and cortical neurons spans a distance greater than any reported primary cortico-cortical connection.

### dLGN-V1 spike synchrony and V1 oscillations

In order to determine whether rhythmicity of synchronous spikes in the thalamocortical pathway were linked to LFP oscillations in V1, we performed a correlation analysis of the synchrony function to different frequencies components in the LFP. This analysis was performed separately for periods where the animals were observing the gray and the checkerboard images. Although spike rates in V1 were lower during gray screen viewing, periods of spontaneously increased spike activity showed rhythmic spike synchrony which was highly correlated with narrowband gamma in V1 (Figure 5A). Similarly, rhythmic spike synchrony that occurred time-locked to the presentation of the checkerboard image was highly correlated with gamma oscillations across a broader frequency band, which overlapped with the evoked broadband gamma, supporting a causal relationship (Figure 5B). Correlation values were statistically significant in the high gamma range during visual stimulation with the checkerboard in 6 of 6 mice (Figure 5D), and during gray screen viewing in 5 of 6 mice (Figure 5C), based on the results of our Monte Carlo hypothesis test.

**Figure 5:**
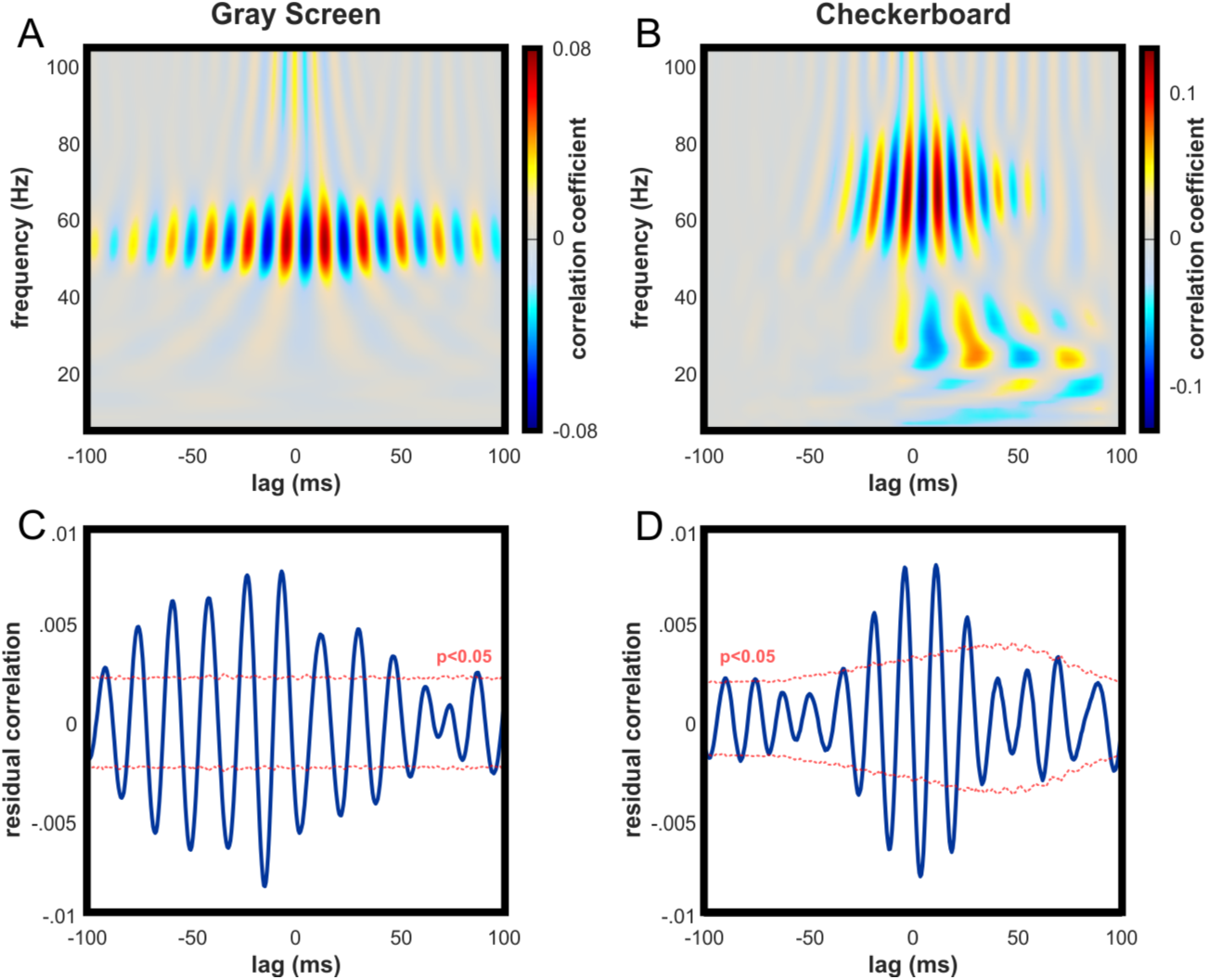
Rhythmic synchronization of spike activity in dLGN and V1 is strongly linked to both narrow and broadband high gamma in V1. A: Example cross-correlation of dLGN-V1 spike synchrony and V1 LFP oscillations across frequency spectra during gray screen viewing. Peak correlation occurs in the narrowband gamma frequency range. B: Same as A for checkerboard viewing. Peak correlation exists across a broader frequency band, consistent with contrast-evoked broadband high gamma in V1. C: Example statistical test for correlation between spike synchrony and bandpass-filtered high gamma in the V1 LFP (50-90Hz) during gray screen viewing. Tested individually, significant peaks and troughs were detected in 5 of 6 animals. 1000 surrogates were generated by jittering dLGN spikes and median surrogate values were subtracted for easier visual interpretation. D: Same as C for checkerboard viewing. Tested individually, significant peaks and troughs were detected in all animals.

## Discussion

We used simultaneous recordings of local field potential and spike activity in six mice to identify consistent patterns of rhythmic interactions between V1 and dLGN associated with neuronal processing of images with changing contrast. Our findings support the hypothesis that both narrowband and broadband high frequency gamma oscillations in V1 are associated with the effective thalamocortical transmission of visual information. The transition from a gray screen image to a high contrast checkerboard image was characterized by a rapid sequence of events: first, bursting of thalamic relay cells in response to contrast onset, followed by the rhythmic occurrence of synchronous spikes in V1 and dLGN in the high gamma range. This spike synchrony was followed by an increase in the amplitude of gamma oscillations in V1, which were phase-locked to gamma oscillations in dLGN. The center frequency of the high gamma response in V1 varied between mice, but had 20-30Hz bandwidth in the frequency range of 50-90Hz. Across stimulus conditions, the amplitude of high gamma in V1 was strongly linked to the degree of phase synchronization with high gamma in dLGN, and the occurrence of synchronous spikes between the regions was significantly correlated with oscillations in the high gamma range of the V1 LFP.

We also observed prominent contrast evoked beta/gamma oscillations in V1 below the narrowband frequency which were not synchronized with activity in dLGN. Recent reports show that oscillations in this frequency range were evoked in V1 by high contrast bars, and that the amplitude of these oscillations corresponded to the contiguous width of the bars (Veit, Hakim et al. 2017). Given that LFP is an aggregate measure of neuronal currents in a recording area (Buzsaki, Anastassiou et al. 2012), this is consistent with the hypothesis that the low gamma we observed in V1 in response to a broad checkerboard image reflects corticocortical interactions independent of rhythmic thalamocortical influences. The evidence that these oscillations were mediated by local somatostatin-expressing interneurons further supports this as an emergent cortical phenomenon.

### Comparison to awake primates

Previous studies have shown that LFP oscillations in separate frequency bands allow for single regions to selectively communicate with multiple others, as if by dedicated “channels” for transmission of sensory information (Colgin, Denninger et al. 2009, Bastos, Briggs et al. 2014). In the context of visual processing in primates, these distinct frequency bands are thought to correspond to mechanisms of top-down versus bottom-up signaling between lower- and higherorder visual cortical regions (Engel, Fries et al. 2001, Richter, Thompson et al. 2017). Interestingly, gamma oscillations first appear in V1, and are driven by the strength of synaptic input from LGN (Ray and Maunsell 2010, Ray and Maunsell 2011), which occurs rhythmically in the beta frequency range (Bastos, Briggs et al.).

Our results suggest that the rhythms that support thalamocortical communication are substantially different in rodents and primates, and that oscillations in the high gamma range are not an emergent phenomenon in the mouse cortex. Previous studies which explored the generation and function of gamma in the mouser retina, dLGN, and V1 strongly support this difference between species (Koepsell, Wang et al. 2009, Saleem, Lien et al. 2017, Storchi, Bedford et al. 2017).

LFP components in the high gamma range are known to arise in macaque visual cortex during increased spiking activity from visual stimulation, and have been studied independently of thalamic activity (Belitski, Gretton et al. 2008, Ray and Maunsell 2011), but there are diverging hypotheses about their role in visual processing. One group determined through an information theoretic analysis that power in the high gamma band (peak at ~74Hz) conveyed the most information about the visual stimulus being presented (Belitski, Gretton et al. 2008). The amplitude of these oscillations clearly co-varied with the visual-evoked spike rate, suggesting a link between high gamma in V1 and excitatory input related to visual information. Such a role for high gamma in monkeys – namely that it is associated with the fidelity of visual information relay to cortex - is consistent with our findings, although the mechanisms remain obscure. In contrast to this interpretation however, another group made the observation that these visual-evoked frequency components are mostly transient (rather than oscillatory) and not band-limited, suggesting that they are most likely a reflection of currents generated by the firing of local units (Ray and Maunsell 2011). Furthermore, high gamma oscillations in the macaque reportedly occupy on the order of 1% of the LFP energy. This is not consistent with our observations in mice, given that oscillations in the high gamma range are visible in raw LFP traces (Figure 1A, C; Figure 2A), and are band-limited (Figure 1B, D-G). Given these considerations in addition to the direct study of simultaneous LGN and V1 recordings in macaque, it seems unlikely that high gamma in V1 is associated with thalamocortical transmission of visual information in primates.

Briefly, in anesthetized cats, rhythmic spike correlations within LGN exist exclusively in the high gamma range (>70Hz), and thalamocortical spike correlations have been observed over broad frequency range (30-100Hz) with a median frequency of approximately 50Hz (Castelo-Branco, Neuenschwander et al. 1998). Rhythmic synchronization was observed between visual cortical neurons at lower frequencies (30-50Hz) (Castelo-Branco, Neuenschwander et al. 1998), consistent with findings in anesthetized rodents (Nase, Singer et al. 2003). This may suggest a role for high gamma synchronization in dedicated thalamocortical communication in cats, but further investigation is necessary. To our knowledge, no studies of simultaneous recordings from LGN and V1 have been conducted in awake cats, making a direct comparison to our findings inappropriate.

### Broad synchrony between thalamic and visual cortical units

The reliability with which we were able to capture significant spike correlations seems worthy of consideration. The fact that all mice exhibited prominent broadband gamma in V1, as well as precise spike correlations between neurons separated by a projection of several millimeters, suggests that broadband gamma driven by stimulus contrast likely plays a role in the synchronization of extensive neuronal populations across these regions. This is consistent with the hypothesized function of thalamocortical circuit architecture, which includes several features that promote synchrony on a broad scale (Jones 2002) and maximize the effectiveness of information relay (Wang, Spencer et al. 2010). These features include stimulus-driven inhibition from the thalamic reticular nucleus, which is activated by both cortical and thalamic neurons and synchronizes spike output from the thalamus (Sherman and Guillery 2002). Furthermore, in mammals such as cats and monkeys, a subpopulation of thalamocortical neurons are broadly-projecting and nonspecific, with the hypothesized role of creating a synchronized matrix across the cortical sensory area (Jones 2001). Similar investigations are still needed to determine the specific cellular mechanisms of thalamocortical synchrony in mice.

## Notes

Conflict of Interest: None

